# C1q/CTRP1 exerts neuroprotective effects in TBI rats by regulating inflammation and autophagy

**DOI:** 10.1101/2020.12.29.424653

**Authors:** Ming Pei, Chaoqun Wang, Zhengdong Li, Jianhua Zhang, Ping Huang, Jiawen Wang, Jiang Huang, Donghua Zou, Yijiu Chen

## Abstract

**Objective:** C1q/CTRP1 is a newly discovered adiponectin protein, which is highly expressed in adipose and heart tissues. Recent studies have revealed that C1q/CTRP1 can regulate metabolism and inhibit inflammation. CTRP1 is also expressed in brain tissues and vascular cells of human and rat, and research on cerebral hemorrhage and cerebral ischemia-reperfusion injury demonstrates that the CTRP family can attenuate secondary brain injury and exert neuroprotective effects. Thus, this study was designed to explore the role of CTRP1 in traumatic brain injury (TBI) and the underlying mechanism.

**Main methods:** Rats were assigned into rCTRP1 group, vehicle group, and sham group. Modified Feeney’s method was used to establish a closed traumatic brain injury model. Morris water maze was used for directional navigation, reverse searching and space exploration tests in rats. In addition, Golgi-Cox staining was utilized to visualize neurons, dendrites and dendritic spines. ELISA was conducted to detect the levels of inflammatory factors (IL-6 and TNF-α). Finally, Western blot was adopted to detect the relative expression of *p*-mTOR and autophagy-related proteins (Beclin-1 and LC3-II).

**Results:** CTRP1 improved the behavioral and histopathological outcomes, inhibited the inflammatory response, activated mTOR and decreased autophagy-associated protein synthesis in TBI rats.

**Conclusion:** CTRP1 exerts neuroprotective effects in TBI rats by regulating inflammation and autophagy and has potential therapeutic properties after TBI.

## Introduction

Traumatic brain injury (TBI) is the main cause of traumatic death and disability globally. The common causes of TBI include violence, traffic accidents, tumble, falls, and sports accidents [1]. TBI is generally divided into primary and secondary brain injury. To be specific, primary TBI is caused by direct external force acting on the central nervous system. Primary TBI often leads to neurometabolic disorders, hippocampal synapse damage, neuron, astrocytic degeneration (CA1/CA3 layer, dentate gyrus) and glutamate excitotoxicity. In addition, with the subsequent destruction of the blood-brain barrier (BBB) and the persistence of neurogenic inflammation [2], there are secondary changes in nerve tissues, including a series of pathophysiological changes, energy metabolism disorders and inflammatory response [3–5], which generally aggravate neuronal necrosis, dendrite and synapse damage [2]. Both primary and secondary TBI can directly cause cognitive and behavioral dysfunction in patients, which seriously affects the patient’s quality of life [6,7]. Therefore, how to attenuate the damage of primary and secondary TBI on the nerve tissue and to enhance the neuroprotective effects has become the present research focus of neuroscience and trauma science.

C1q/TNF-related protein-1 (CTRP1) is a protein cytokine containing 281 amino acids, which belongs to the CTRP family. And the CTRP family is mostly involved in regulating inflammation and metabolism [8]. Similar to other CTRP family members, CTRP1 is expressed in adipose and heart tissues [9,10]. CTRP1-deficient mice show increased myocardial infarction area caused by ischemia reperfusion injury (IRI), cardiomyocyte apoptosis and expression of pro-inflammatory genes, while the up-regulation of CTRP1 protein expression can attenuate myocardial injury, indicating that CTRP1 can regulate cardiomyocyte metabolism, inhibit inflammatory response and protect the damaged cardiomyocytes [10]. Additionally, inflammatory response and pro-inflammatory cytokines can increase the secretion of CTRP1 [11–13]. Relevant studies have shown that the increased CTRP1 protein expression can suppress the inflammatory response caused by cerebral IRI [14]. Moreover, CTRP1 can regulate the autophagy of glial cells by activating the Akt/mTOR signaling pathway, thereby exerting neuroprotective effects [14]. Based on the above research, this study aimed to confirm the effectiveness of CTRP1 on neuroprotection, and to investigate its effect on memory, cognitive and behavioral functions in TBI rats. In addition, we further revealed the mechanism of the neuroprotective effects of CTRP1 in TBI rats.

## 1. Materials and methods

### 1.1 Animals

The use of animals and the animal procedures were conducted following the guidelines approved and formulated by the Animal Care and Use Committee of the Xuzhou Medical University. A total of 80 healthy male specific pathogen free (SPF) Sprague-Dawley (SD) rats (10 to 12 weeks old, 220 to 250 g of weight) were purchased from the Animal Experimental Center of Xuzhou Medical University. Rats were maintained under SPF condition, with the temperature of 25±1 °C, the relative humidity of 40%-60% and the light/dark cycle of 12 h/12 h. Rats were freely accessible to food and water. The rats were adaptively fed for one week before the experiment. All surgery was performed under sodium pentobarbital anesthesia, and all efforts were made to minimize suffering.

### 1.2 Construction of the TBI model

The modified Feeney’s method was used to establish the closed TBI model^16^. In brief, rats were fasted and deprived of water for 12 h before operation. The head was fixed on the stereotaxic apparatus, followed by shearing and disinfection of the operation area. Afterwards, the scalp was cut along the midline sagittal of the head to expose the right parietal skull. To penetrate the skull and open a circular window 5 mm in diameter while preventing endocranial injury, a dental drill was used on the skull at a distance of 3.5 mm to the right of the skull midline and about 0.2 mm from the bregma. Subsequently, a 40 g skull batting stick was released from a height of 20 cm of the stereotaxic apparatus vertically fixed to the cannula, so as to control the subsidence depth of 0.2 cm and the diameter of striking end of approximately 0.2 cm, causing contusion of the right hemisphere. Finally, the skull defect was sealed with bone wax, and the scalp was intermittently sutured. Rats in the sham group were subjected to skull window exposure without further hitting. Symptoms such as limb twitch, urinary incontinence, nasal bleeding and a few seconds of apnea were shown in rats after the hit. Based on the neurological severity score (NSS) at 0.5 h and 1 h after injury, the rats with common TBI were screened, excluding eight rats that died or did not present moderate TBI.

### 1.3 Grouping and drug administration

The remaining 72 rats were randomly grouped into three categories matching the NSS results, including the rCTRP1 group (TBI + rCTRP1 recombinant protein), which was given an acute intracerebroventricular injection of 80 μg/kg mouse-derived rCTRP1 recombinant protein (ANNORON, China) every 24 h for up to 7d from half an hour after TBI; the vehicle group (TBI +vehicle), which was administered with the same amount of normal saline at equal frequency to that of the rCTRP1 group after craniocerebral trauma; and the sham group, which received a craniotomy without TBI. Three time points after TBI, namely, 24 h, 72 h and 1 w, were chosen, and eight rats in each group were examined at each time point.

### 1.4 Morris water maze

The Morris water maze consisted of a round dark metallic pool 160 cm in diameter and 60 cm in depth that was filled with water (22±0.5°C). Water was made opaque by the addition of a dark nontoxic water-based paint to a depth of 50 cm and surrounded by a dark curtain. The pool was virtually divided into four quadrants. An escape platform (12 ×12 cm) was submerged at 1 cm below the opaque water surface and located in the center of one quadrant of the maze, approximately 30 cm from the edge of pool. Randomized to one of the four quadrants, rats were allowed to search for the hidden platform for 1 min. The rats were placed on the platform manually for 15 s if they exceeded the allotted time. Rats were trained for five consecutive days before the experiment. Then, at 24 h, 72 h and 1 w after TBI, rats were allowed to seek the platform in the 1-min test. After the water maze, rats were sacrificed. Hippocampal tissue samples were immediately removed and stored at −80°C.

### 1.5 Golgi-Cox staining and microscopy procedures

One side of the hippocampus of rats in each group was transferred into the impregnation solution at room temperature based on the instructions of the Golgi-Cox OptimStain Prekit (HiTO, USA). The impregnation solution was placed and stored at room temperature in dark for two weeks. The tissue was subsequently transferred into solution-3 for 12 h at 4°C. Then solution-3 was replaced and stored for 48 h at 4°C. The temperature of isopentane was attenuated to −70 °C using dry ice. The hippocampal tissue was then immersed in the isopentane and cooled down for approximately 40 s, and the absorbent paper was used to remove excess isopentane from the tissue. The freezing microtome was pre-cooled to −19°C, which was used to slowly cut the tissue into sections 120-μm in thickness. The sections were transferred to a gelatin-coated slide and dried at room temperature in darkness overnight. Thereafter, the slides were rinsed in distilled water twice for 3 min each time. Later, the slides were placed in the staining mixing solution for 10 min and then in renewed distilled water twice for 4 min each time. The slides were then dehydrated in 50%, 75% and 95% ethanol (5 min each) and in 100% ethanol thrice (5 min each). Afterwards, the slides were cleaned in xylene twice (5 min each), sealed with a coverslip, and viewed by light microscopy. At least 10 neurons were randomly selected in each slice (magnification 200 times) by a trained observer blind to the experimental condition. Equi-distant (10 μm) concentric rings were placed over the tracings of the dendritic tree. The total dendritic arborization and dendritic length were measured by the amount of ring intersections with the dendritic tree. Over 10 primary dendritic branches at a length of ≥20μm were traced (at 1000×). The amount of dendritic spine was computed utilizing the analysis system of Image pro-plus 6.0.

### 1.6 Detection of IL-6 and TNF-α by ELISA

Hippocampal tissue was harvested immediately and homogenized in lysis buffer, followed by centrifugation at 8000g for 10 min at 4°C to collect the supernatants. The levels of IL-6 and TNF-α were determined by ELISA kit purchased from Abcam Company (ab100785 and ab100772, Abcam, America) according to the manufacturer’s instruction.

### 1.7 Western blot analysis

Hippocampal tissue was centrifuged at 12000 rpm for 15 min and homogenized in ice-cold tissue lysis buffer (50 mm Tris, PH 7.5, 0.15 mm NaCl, 2% NP-40, 0.5% sodium deoxycholate, 4% SDS, and protease and phosphatase inhibitor cocktails) for 15 min. In every supernatant fraction, the BAC protein assay kit was used to measure the total protein concentration (Pierce, Rockford, IL, USA), and 15 μg Beclin-1, LC3-II and *p*-mTOR were separated by electrophoresis and then transferred onto the nitrocellulose membranes (Bio-Rad; Trans-Blot Turbo Transfer System). After several washes with TBST buffer, the membranes were blocked for 2 h with the blocking buffer (LI-COR Biosciences, Lincoln, Nebraska, USA) and later incubated with rabbit anti-Beclin-1 (1:4000; Abcam, America), rabbit anti-LC3-II (1:2000; Abcam, America), polyclonal rabbit Anti-mTOR (1:5000, Abcam, America), and rabbit anti-GAPDH (1:10,000; Abcam, America) in TBST at 4°C overnight. After four washes, the HRP-linked secondary antibodies (1:5,000, Boster Bioengineering, China) were incubated in dark for 1 h. The Image J software was used to quantify the bands (NIH, Bethesda, MD, USA).

### 1.8 Statistical analysis

The description of continuous variables in our dataset was expressed as mean ± standard deviation. The repeated measurement ANOVA was employed to analyze the behavioral statistics during the training period. The one-way ANOVA was utilized to examine the spine density, dendritic length and branching, as well as the ELISA and Western blot data. Further multiple comparisons were undertaken via the LSD method if the overall ANOVA analysis was significant. All the statistical analyses in the present study were implemented by the SPSS software (version 19.0 for Windows, SPSS Inc., USA) and the significance level was set to be 0.05 throughout our study.

## 2 Results

### 2.1 The expression of CTRP1 in hippocampus

The expression of CTRP1 in rat hippocampus was detected at 24 h after TBI. The results showed that the expression of CTRP1 increased in rCTRP1 group and vehicle group compared with sham group (p <0.001). In addition, the expression of CTRP1 was significantly higher in rCTRP1 group than that in vehicle group (p<0.001) (Fig 1A B).

**Fig 1.**
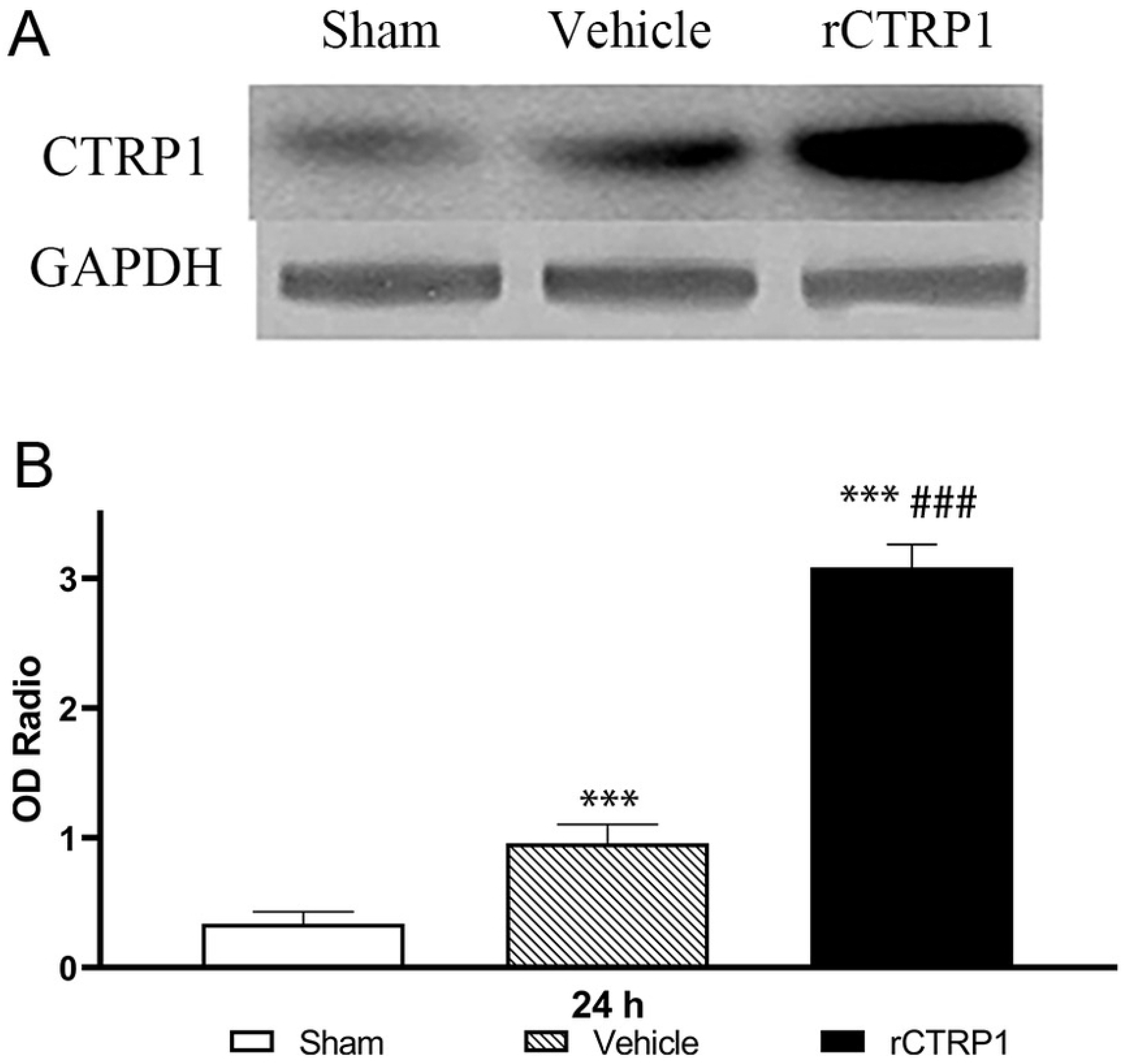
Expression of CTRP1 in hippocampus at 24h after TBI. *** p < 0.001 vs. sham group, ###p < 0.001 vs vehicle group.

### 2.2 Results of memory and spatial learning

#### 2.2.1 Spatial orientation ability

The escape latency is defined as the time from entering water to boarding the platform of rats, which is one of the commonly indicators of spatial orientation ability. In this study, the escape latency time was detected in rats of three different groups at 24 h, 72 h, and 1 w after craniocerebral trauma. As a result, the escape latency was significantly longer in rCTRP1 group and vehicle group than that in sham group at the first two time points (p<0.005), and there was no significant difference in the escape latency between rCTRP1 group and sham group at 1 w after craniocerebral trauma. However, the time was significantly shorter in rCTRP1 group than that in vehicle group at 1 w after TBI (p<0.05) (Fig 2A).

**Fig 2.**
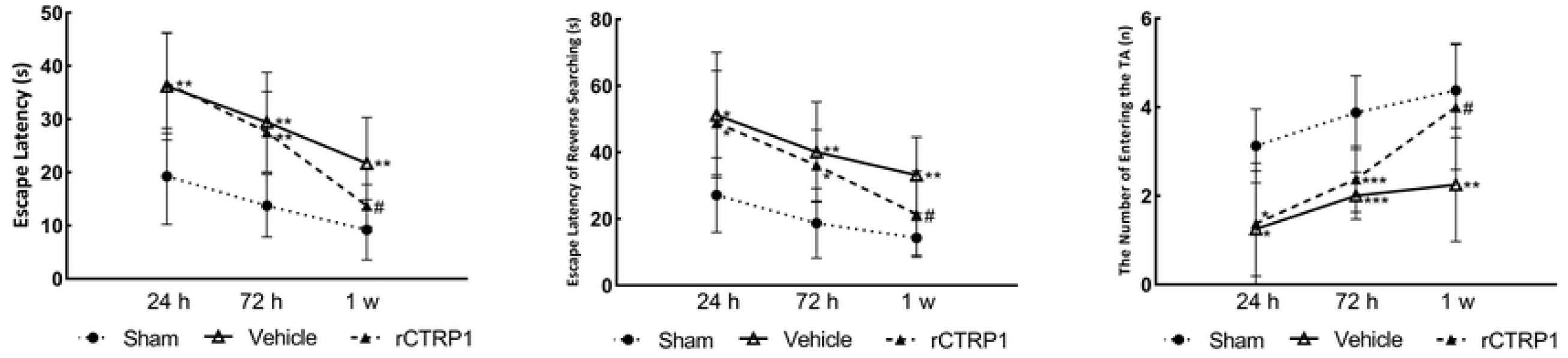
The results of two Morris water maze tests in three groups at 24h, 72h and 1w after TBI. **(A)** Escape latency to locate the target platform. **(B)** Escape latency of reverse searching to locate the target platform. **(C)** After the platform was removed, the number of entries into the target quadrant (TA). *p < 0.05 vs. sham group, ** p < 0.005 vs. sham group, ***p<0.001 vs. sham group, # p < 0.05 vs vehicle group.

#### 2.2.2 Reverse searching ability

The second quadrant of the platform was changed to the fourth quadrant, followed by re-examination of the escape latency of rats, which was used as an indicator of working memory ability. The escape latency of reserve searching was significantly prolonged in rCTRP1 group and vehicle group than that in sham group at 24 h and 72 h after craniocerebral trauma (p<0.05), and there was no significant difference in the time between rCTRP1 group and sham group at 1 w after craniocerebral trauma. However, the time was significantly shorter in rCTRP1 group than that in vehicle group at 1 w after TBI (p<0.05) (Fig 2B).

#### 2.2.3 Spatial exploration ability

The number of entries into the target quadrant (TA) after removing the underwater platform is one of the commonly used indicators to detect the spatial exploration ability of rats. In this study, the number of entries into the TA was significantly less in rCTRP1 group and vehicle group than that in sham group at 24 h and 72 h after TBI (p<0.05), and there was no significant difference in the number between rCTRP1 group and sham group at 1 w after craniocerebral trauma. While the number significantly increased in rCTRP1 group compared to vehicle group at 1 w after TBI (p<0.05) (Fig 2C).

### 2.3 Changes in dendrites and dendritic spines

The alterations in dendrites and dendritic spines were observed at 24 h, 72 h and 1 w after TBI, respectively. The length and branches of dendrites significantly decreased in rCTRP1 group and vehicle group than those in sham group at 72 h and 1 w after TBI (p<0.05), and there was no significant difference in those between rCTRP1 group and sham group at 24 h after craniocerebral trauma. Moreover, the length and branches of dendrites significantly increased in rCTRP1 group compared to those in vehicle group at 1 w after TBI (p<0.05) (Fig 3ABC, Fig 4A). The dendritic spines decreased in rCTRP1 group and vehicle group than those in sham group at 24 h and 72 h after TBI (p<0.005), and there was no significant difference in those between rCTRP1 group and sham group at 1 w after craniocerebral trauma. However, the dendritic spines significantly increased in rCTRP1 group compared to those in vehicle group at 72 h and 1 w after TBI (p<0.05) (Fig 3 DEF, Fig 4B).

**Fig 3.**
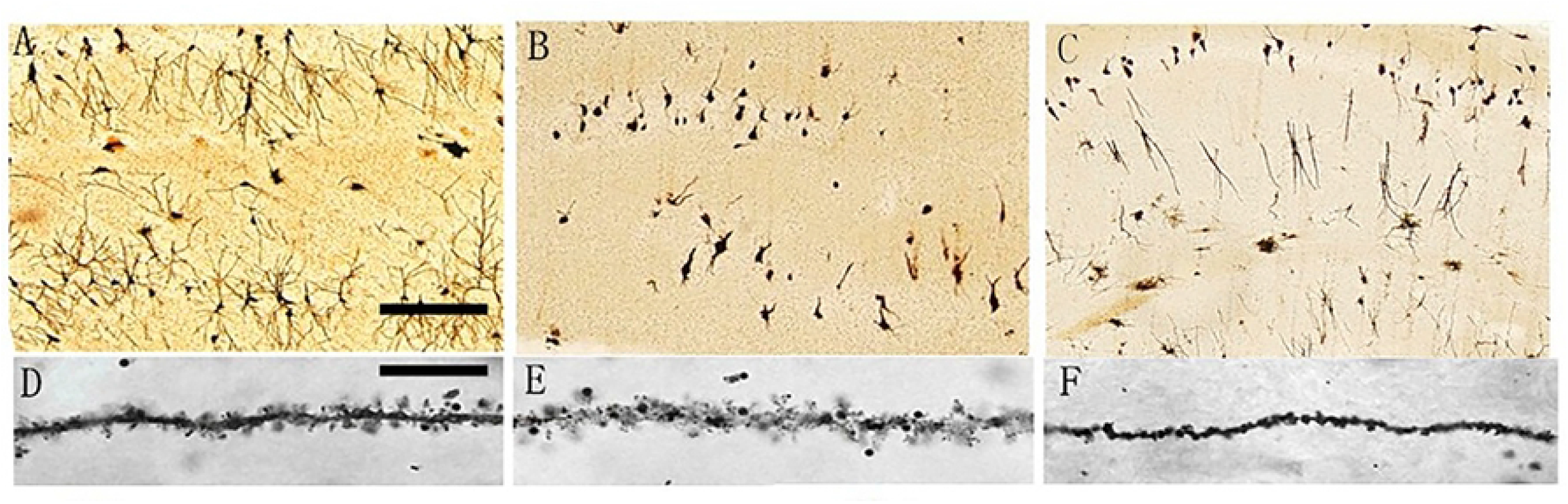
Changes in dendrites and dendritic spines at 72h after TBI. **(A)** (the black bar is 50 μm) and **(D)** (the black bar is 10 μm), sham group; **(B)** and **(E)**, vehicle group; **(C)** and **(F)**, rCTRP1 group.

**Fig 4.**
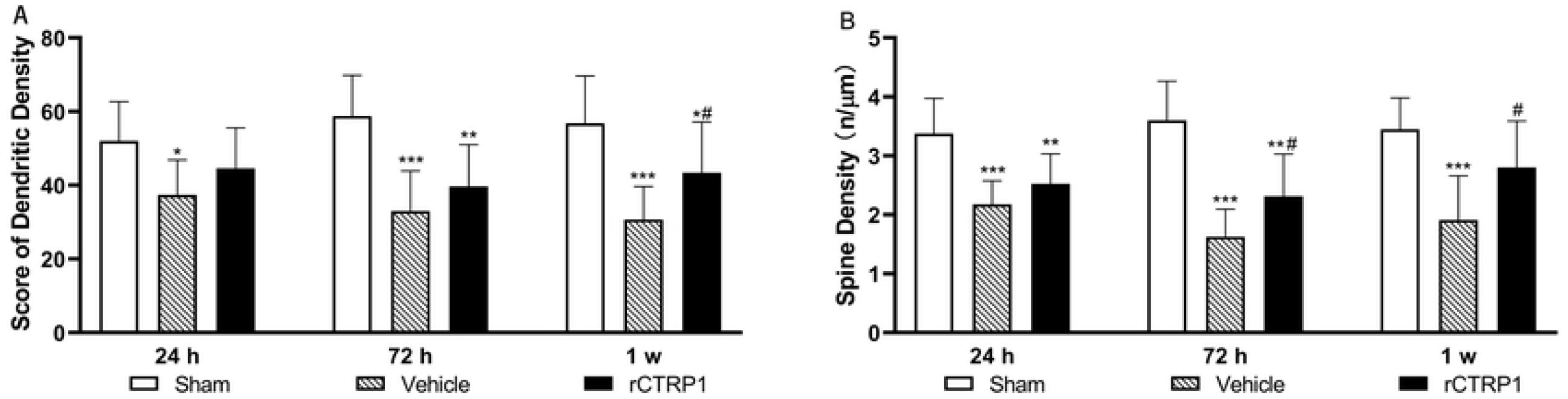
The histogram of changes in dendrites and dendritic spines at 24h, 72h and 1w after TBI. **(A)** The number of ring intersections of the dendritic arborization. **(B)** The density of the dendritic spines. * p < 0.05 vs. sham group, **p<0.005 vs. sham group, *** p<0.001 vs. sham group, # p < 0.05 vs vehicle group.

### 2.4 Effects of CTRP1 protein on neuroinflammation

As presented in Fig. 5, after TBI, enhanced concentrations of TNF-α in rCTRP1 group and vehicle group were observed at 24 h, 72 h and 1 w after TBI compared to sham group (p < 0.05); typically, the concentrations rapidly increased within 24 h after TBI and then slowly recovered to normal levels. The change trend of IL-6 concentrations in the three groups was similar to that of TNF-α at the three time points. The IL-6 and TNF-α levels in rCTRP1 group significantly decreased compared to those in vehicle group at 24 h and 72 h after TBI (p<0.001). The results suggested that the inflammatory response was relatively significant within 1 w after craniocerebral trauma and CTRP1 exerted anti-inflammatory effects in the early stage of injury.

**Fig 5.**
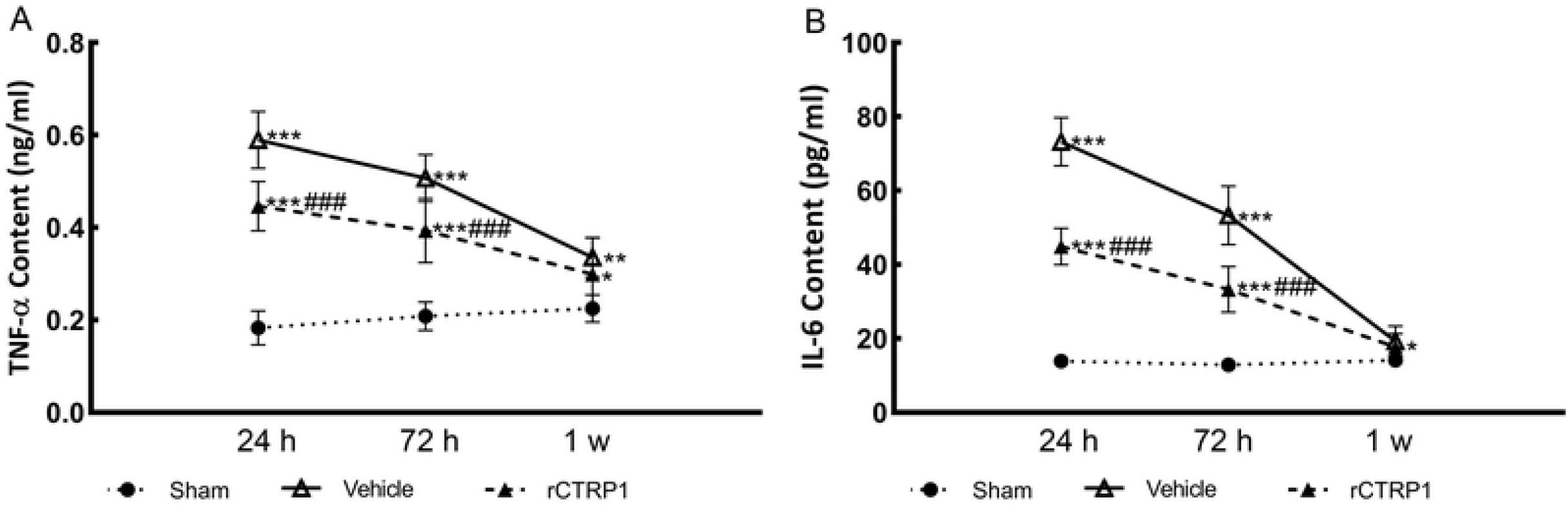
The changes of TNF-α and IL-6 levers in each group at different time points after craniocerebral trauma. * p < 0.05 vs. sham group, ** p < 0.005 vs. sham group, *** p < 0.001 vs. sham group, ### p < 0.001 vs vehicle group.

### 2.5 Effects of CTRP1 protein on autophagy

Western blot was used to detect the expression of Beclin-1, LC3-II and *p*-mTOR in the hippocampus at 24 h, 72 h and 7 d after TBI and to demonstrate the changes in autophagy after TBI and the regulatory roles of CTRP1 on autophagy (Fig 6ABCD). The expression of the autophagy-related proteins Beclin-1 and LC3 - II in hippocampus of vehicle group was significantly higher than that in sham group at 24 h, 72 h, and 1 w after craniocerebral trauma (p<0.05). The properly increased level of autophagy can attenuate cellular functions and decrease energy consumption, which is one of the self-protection mechanisms of damaged cells. However, intense autophagy can also cause cell death. The expression of Beclin-1 and LC3-II in hippocampus tissue in rCTRP1 group was significantly lower than that in vehicle group at three time points (p<0.001), indicating that CTRP1 attenuated autophagic injury after craniocerebral trauma. The expression of *p*-mTOR was significantly lower in vehicle group than that in sham group at 24 h, 72 h and 7 d after TBI (p<0.001). Additionally, the expression of *p*-mTOR was significantly higher in rCTRP1 group than that in vehicle group at three time points (p<0.001), suggesting that the CTRP1 recombinant protein was involved in the regulation of the mTOR pathway.

**Fig 6.**
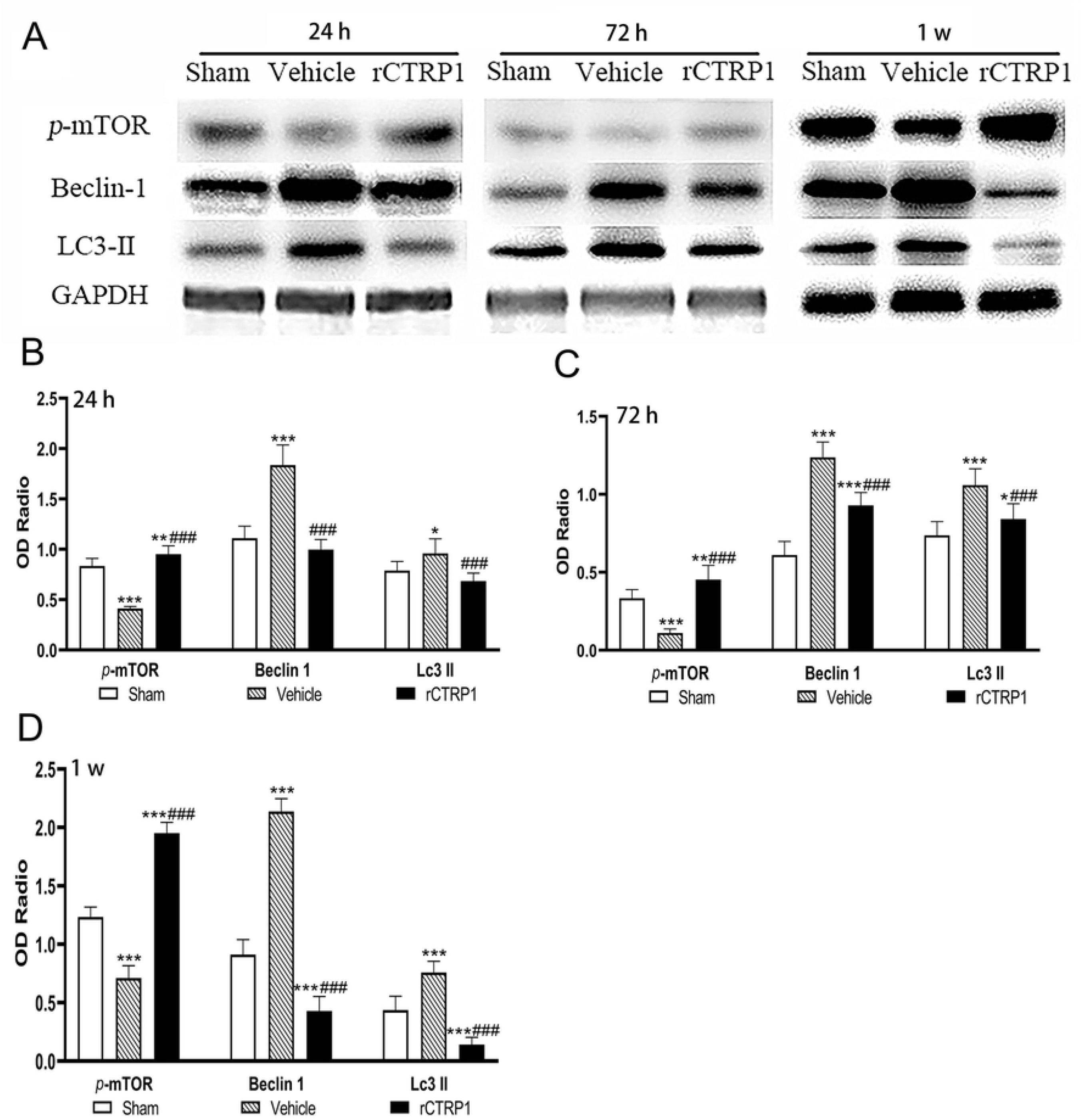
Western-blot analysis was used to detect the expression of *p*-mTOR, Beclin-1 and LC3-II in the hippocampus of each group after craniocerebral trauma. **(A)** Relative expression of *p*-mTOR, Beclin-1 and LC3-II in each group at three time points. **(B-D)** The histogram of the expression of *p-* mTOR, Beclin-1 and LC3-II in each group at 24h, 72h and 1w after craniocerebral trauma. *p < 0.05 vs. sham group, **p < 0.005 vs. sham group, ***p < 0.001 vs. sham group, ###p < 0.001 vs vehicle group.

## 3 Discussion

CTRP1 is a secretory glycoprotein that is expressed in the heart, liver, kidney, placenta, brain and interstitial vascular cells (including macrophages) in human and rat [15–18]. Previous studies have shown that the level of circulating CTRP1 elevates in patients with type 2 diabetes [19–20], coronary artery disease [21] and hypertension [22], and CTRP1 may be involved in the vasculitis and coagulation process in the acute phase of Kawasaki disease [23]. Studies have also demonstrated that CTRP1 possesses insulin-sensitizing effects [24,25]. Recent studies have revealed that the expression of CTRP1 significantly increases in the serum of stroke patients, and is positively correlated with the high-sensitivity C-reactive protein, suggesting that CTRP1 may exert a neuroprotective effect after ischemic stroke [14]. In our study, we found that the expression of CTRP1 increased in the hippocampus of rats at 24 h after craniocerebral trauma and CTRP1 improved the behavioral and histopathological outcomes, indicating that CTRP1 might exert a neuroprotective effect after craniocerebral trauma, which was consistent with the above studies concerning stroke.

A variety of studies have reported that CTPR1 can inhibit inflammation. For instance, CTPR1 protects the heart from IRI by attenuating myocardial cell apoptosis and inflammation [26]. CTRP1 is associated with ischemic heart disease. The myocardial infarction area, cardiomyocyte apoptosis and pro-inflammatory gene expression following IRI are up-regulated in CTPR1 knockout mice, compared with wild-type (WT) mice. In contrast, the up-regulation of CTRP1 protein can attenuate myocardial damage after IRI in WT mice. Treatment of cardiomyocytes with CTRP1 can reduce the hypoxia-reoxygenation-induced apoptosis and the LSP-stimulated expression of proinflammatory cytokines, which can be reversed by inhibiting the sphingosine-1-phosphate (S1P) signaling [26]. Therefore, CTRP1 can be used as an endogenous cardioprotective factor with anti-apoptotic and anti-inflammatory effects. In addition, the detection of CTRP1 expression in the plasma of patients with different types of stroke reveals that the expression level of CTRP1 in atherosclerosis-related strokes is significantly lower than that in other types of strokes, indicating that CTRP1 may be involved in the occurrence and development of stroke via atherosclerosis and the mechanism is associated with the suppression of inflammatory response of macrophages, thereby protecting brain function [27]. In our study, we found that rCTRP1 decreased the expression of inflammatory factors (TNF-α and IL-6) in the hippocampus of rats after TBI.

Autophagy exerts a dual role in TBI. On the one hand, enhanced autophagy can remove the damaged proteins and organelles, and reduce mitochondrial energy consumption, which are beneficial to maintain cell homeostasis. On the other hand, the enhanced autophagy can simultaneously destroy functional macromolecular substances and organelles in cells to aggravate cell damage [28]. Autophagy is regulated by a variety of molecular mechanisms, and the Akt/mTOR signaling pathway plays an important role in regulating autophagy [29]. In addition, the Akt/mTOR pathway attenuates neuronal apoptosis by suppressing calcification-dependent pathways, which has a protective effect in neurological diseases [30,31]. CTRP1 can specifically activate the Akt (protein kinase B) [32] and MAPK signaling pathways in myotubes of the differentiated mice [25]. Huilin Wang *et al.* [14] found that activation of the Akt/mTOR pathway by IGF-1 significantly inhibited the promoting role of si-CTRP1 in microglial autophagy, while A6730 inhibited the Akt and reversed the inhibitory effect of CTRP1 recombinant protein on microglial autophagy. Consistent with previous reports, we discovered that rCTRP1 reduced the expression of hippocampal autophagy-related proteins LC3-II and Beclin-1 by activating the mTOR signaling pathway in rats, and exerted an inhibitory effect on autophagy in hippocampus tissue after TBI, thereby decreasing autophagic injury caused by craniocerebral trauma. These findings suggest that rCTRP1 plays a neuroprotective effect by activating the mTOR signaling pathway.

## 4. Conclusion

In this study, we find that the CTRP1 recombinant protein can improve the behavioral and histopathological outcomes, inhibit inflammatory response, activate mTOR and decrease autophagy-associated protein synthesis in TBI rats. Therefore, CTRP1 exerts neuroprotective effects in TBI rats by regulating inflammation and autophagy and has potential therapeutic properties after TBI.

## Supporting information

S1 Dataset. Raw data.

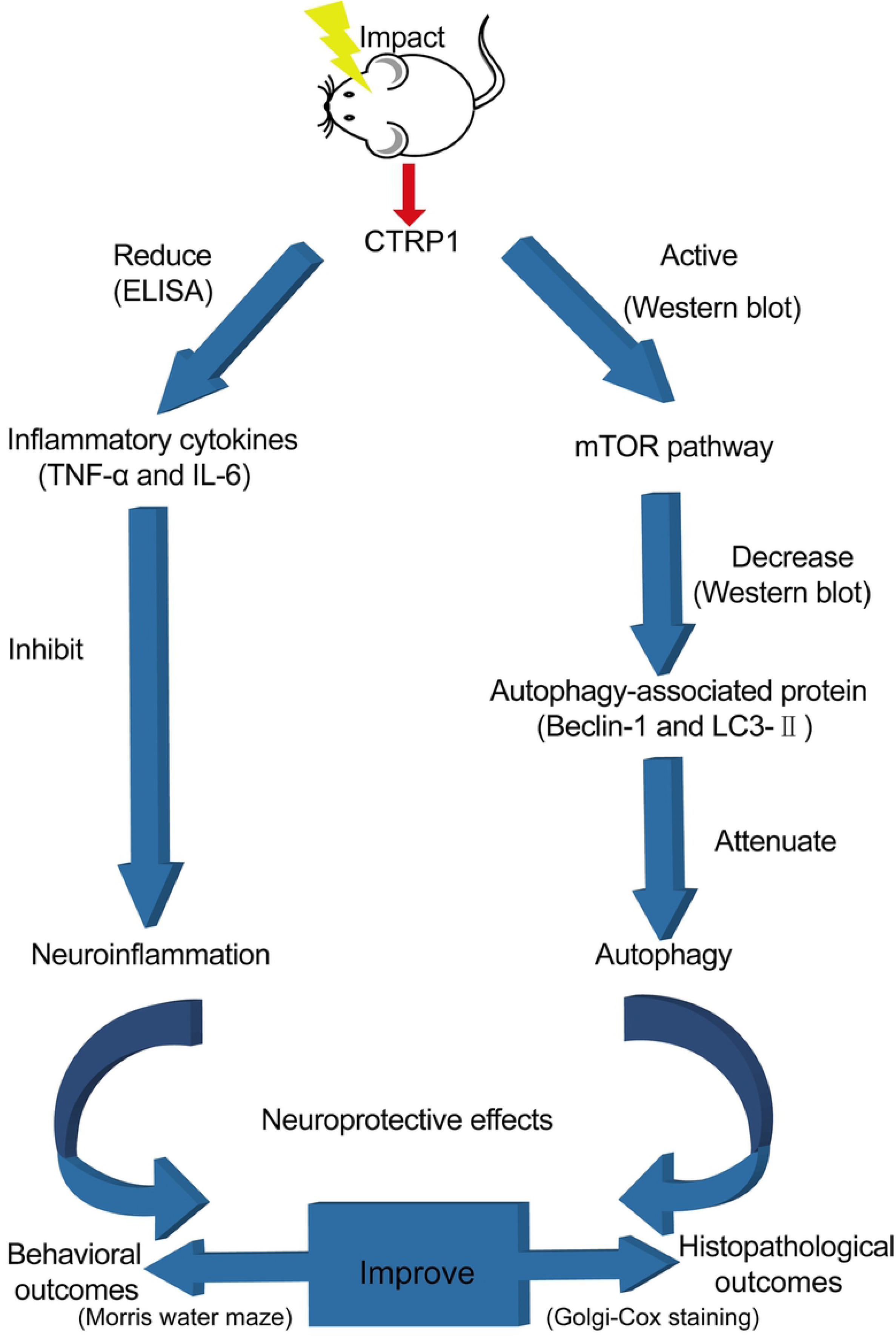

